# Size and mass prediction of almond kernels using machine learning image processing

**DOI:** 10.1101/736348

**Authors:** Sriram K Vidyarthi, Rakhee Tiwari, Samrendra K Singh

## Abstract

After harvesting almond crop, accurate measurement of almond kernel sizes is a significant specification to plan, develop and enhance almond processing operations. The size and mass of the individual almond kernels are vital parameters usually associated with almond quality, particularly head almond yield. In this study, we propose a novel methodology that combines image processing and machine-learning ensemble that accurately measures the size and mass of whole raw almond kernels (classification - Nonpareil) simultaneously. We have developed an image-processing algorithm using recursive method to identify the individual almond kernels from an image and estimate the size of the kernels based on the occupied pixels by a kernel. The number of pixels representing an almond kernel was used as its digital fingerprint to predict its size and mass. Various popular machine learning (ML) models were implemented to build a stacked ensemble model (SEM), predicting the mass of the individual almond kernels based on the features derived from the pixels of the individual kernels in the image. The prediction accuracy and robustness of image processing and SEM were analyzed using uncertainty quantification. The mean error in estimating the average length of 1000 almond kernel was 3.12%. Similarly, mean errors associated with predicting the 1000 kernel mass were 0.63%. The developed algorithm in almond imaging in this study can be used to facilitate a rapid almond yield and quality appraisals.

## 1 Introduction

Almond (*Prunus dulcis*) is one of the finest agricultural products in the world that provides various health benefits, making it one of the healthiest snacks.

Internationally, the United States remains the largest producer of almonds in the world, which harvests over 80% of the world crop supply. California is the largest almond producer in the United States where almonds are grown commercially. In 2018, the total annual production of almond in California was about 2.26 billion lbs at a value of about $5.7 billion (ABC, 2018; AgMRC, 2018; NASS-USDA, 2018). There are 49 varieties of almonds grown in the United States. Four of these varieties, namely Nonpareil, Carmel, California types, and Mission types account for about 85% of all almonds grown in California (NFF, 2019). About 31% of the crop is consumed within the U.S and the rest is exported worldwide. Other than the U.S, the Middle East and European countries are the major producers of almonds in the world. Germany is the largest importer of almonds followed by Japan and the Netherlands.

In the U.S, the almonds are mainly consumed as snacks and the ingredients in the processed food, such as cereal and granola bars. In addition, almonds are used in non-dairy alternative, such as almond milk, offering a low-fat and high protein option for consumers. Other uses of almonds are in home baking, almond paste, almond oil or the blend of almond and other oil, gluten-free flours, ice-creams, food service outlets and confectionaries (AgMRC, 2018; Altan et al., 2011; Debnath et al., 2010; Mange et al., 1987; Rumsey and McCarthy, 2012).

### 1.1 Importance and methods of determining the size and mass of almond kernels

The size of almond kernel is an important attribute, which significantly helps in defining its class internationally. Almonds are commonly sorted and sold in four grades based on USDA standards of kernel quality, which is defined by the amount of chips and scratches, foreign material, and splits and broken almonds. The almonds are graded into four categories. These are Extra No. 1, No. 1 Supreme, U.S. Select Sheller Run, and U.S. Standard Sheller Run, which are described in Table 1. Furthermore, processing of almonds takes place based on kernel sizing and its forms, such as blanched and natural almonds. Additionally, sorting of almonds is done based on the grade and size which helps define the number of almond kernels per ounce. For example, an almond size of ‘18/20’ is the largest almond size which denotes the yields of 18-20 almonds per ounce. Similarly, ‘36/40’ is generally the smallest sized almond which yields between 36-40 almonds per ounce (AgMRC, 2018; Nuragro, 2017; USDA, 2018).

**Table 1:** USDA grading of whole almond kernels (USDA, 2018)

| Almond grading | Quality | Chips and Scratches | Split and broken | Foreign materials |
| --- | --- | --- | --- | --- |
| U.S. Fancy | Highest | < 5% | < 1% | < 0.05% |
| U.S. Extra No. 1 | Highest | < 5% | < 1% | < 0.05% |
| U.S. No. 1 (Supreme) | 2nd grade | < 10% | < 1% | < 0.05% |
| U.S. Select Sheller Run (SSR) | 3rd grade | < 20% | < 5% | < 0.1% |
| U.S. Standard Sheller Run | 4th grade | < 35% | < 15% | < 0.2% |

Almond kernel size is also one of the major indicators of almond quality. The length, chips, scratches and split in almond kernels provide consistency in the almond samples, which is an important aspect to customers. The importance of maintaining kernel size is significant for the almond industry because of the various forms in which the product is consumed, including whole, halves, slices and pieces. The market value of the almond is different based on the kernel length. Larger average kernel size may contribute to a higher price yield. For example, a larger kernel size of almond is considered of supreme quality and typically has higher price. Similarly, broken almonds with lengths that are less than three-fourths of the unbroken kernel length (including visual characteristics as chips, scratches and split kernel) have relatively lower market values (AMS-USDA, 2018).

Adjusted kernel weight has been a conventional computation method to compute the almond kernel weight which includes a calculation of percentages for specified measurements of edible kernels, inedible kernels, foreign materials, and excess moisture (AMS-USDA, 2018). This computational method is tedious, slow and less accurate due to involvement of the manual nature of measurements and their reporting techniques cannot be very effective when various visual aspects of the kernel, such as chips, scratches and splits are considered.

The conventional adjusted kernel weight was later replaced by a computer-based database program which computed and totaled the percentages for the specified measurements and made minor automatic rounding adjustments, when needed, to equate the total to 100%. However, USDA discovered that the program occasionally made some rounding error at the nearest hundredth decimal. Therefore, the inspection service began testing a new web-based program in 2017 aiming replacement of the computer-based program with more accuracy up to the nearest thousandth decimal rounding (AMS-USDA, 2018).

Based on the recent development, kernel size measurement can be performed using a very effective technique called image analysis. This technique helps achieve accurate and effective results faster with relatively higher throughput. Image processing technologies can be useful to classify the almond kernels based on digital imaging of almond kernels, computer scanning and analysis of kernels using imaging software by calculating the kernel areas and pixel values. For instance, SHAPE is a technique that generates elliptic Fourier descriptors of 2- and 3-dimensional characteristics to evaluate shape from pictures of horizontally and vertically oriented kernel. This method requires preparation of each kernel, making it labor-intensive (Williams et al., 2013). SeedCount (Next Instruments, NSW, Australia) evaluates the variation and sample mean specifically with help of image analysis to measure the size of kernels in a sample. However, this technique is tedious and not economical, particularly for large number of samples (Whan et al., 2014).

Whan et al. (2014) implemented another technique called GrainScan to analyze the measurements and color of wheat and Brachypodium distachyon seeds, which utilized reflected light to capture color information from color space (CIELAB).

ImageJ is a Java based global image analysis software that can be implemented on a variety of grains to analyze various kernel parameters, such as seed shape and size in a range, including wheat and rice (Li et al., 2009). Smartgrain is another image analysis system which helps in analyzing the seed characteristics based on area, perimeter, width, and length by constructing ellipses on identified kernels (Tanabata et al., 2012).

However, high-throughput measurement to determine kernel size is still very challenging with all these current techniques as these methods are still in development stage.

### 1.2 Machine learning in almond research

For the first time, machine learning (ML) technology was implemented for almond measurements in this study using stacked ensemble model (SEM). A predictive model system can be designed in the form of a known data set by implementing Machine-learning model a (Bejagam et al., 2018).

Mirzabe et al. (2013) measured the dimensions of almond kernels and then compared their physical properties, such as bulk density, true density, porosity and coefficient of friction using various statistical regression models, such as linear and quadratic regression analysis and distribution methods, including Normal distribution, Weibull distribution and Log-normal distribution which can be a tedious process. They found no correlation between dimensions of kernels and dimensions of nuts, however, increase in nuts’ length resulted in longer kernels for most of the nuts.

Teimouri et al. (2014) presented an image processing method combined with machine learning to analyze and evaluate segmentation of almond images with different classes, such as normal almond, broken and split almond, shell of almond, wrinkled almond and doubles or twins of almond. They implemented artificial neural networks (ANNs) to classify the images into three categories – objects, shadow and background. They compared the optimum ANN model results with Otsu, dynamic thresholding and watershed methods. They performed sensitivity analysis and obtained high mean values of sensitivity, specificity and accuracy for detected almonds from images.

Halac et al (2017) implemented various supervised machine learning methods, specifically multi-class support vector machines and artificial neural networks to classify different type of almonds. Their method relied on the principal component analysis to identify the most significant shape and color parameters. Their study revealed that the use of support vector machine attained higher levels of accuracy in almond classification than artificial neural networks. Although the results showed higher accuracy overall, the models had increased levels of computational time and complexity.

The conventional techniques of segregating almond kernels’ mass and dimension can have several constrains with respect to time and reliability. This process can be even more complex when performed on a large scale. Hence, we have developed a unique and robust method that accurately identifies the almond kernels and measures their dimensional properties, including individual mass and dimension. Our procedure includes image processing of almond kernel samples to identify the individual almond kernels and then use stacked ensembled model (SEM) image processing to predict their mass. We employed bootstrapped sampling method and performed uncertainty quantification (UQ) to assess the accuracy and robustness of the ML methods. Using the bootstrapping technique, we implemented a unique method with original data set of size n to create NBS number of data sets, each of size n, as described by Singh et al (2019). The objectives of this paper are to accurately predict the size and mass of the almond kernels by applying the image processing in conjunction with the SEM. In addition, we quantified the errors and uncertainties in our model predictions.

## 2 Materials and methods

### 2.1 Almond samples

California-grown whole raw almond kernels (classification - Nonpareil) were purchased from RPAC, LLC, Los Banos, California, USA for conducting the tests in this study. The almonds were harvested in 2018 season. After harvesting, as per the USDA guidelines, almond kernels were processed and stored in the controlled environment required for improved shelf life. After the purchase of almond kernels, the almond samples were procured in Ziploc^®^ bags and stored in the room temperature.

### 2.2 Size and weight measurement of almond kernels

100 almond kernels were manually spread on a black color surface divided into grids of 10 rows x 10 columns. Each kernel was placed in a separate grid. The lengths and breadths of each kernel were measured using a Vernier caliper (Pittsburgh® 6” Digital Caliper; Model - HFT47257; Camarillo, California, USA) with an accuracy and resolution of 0.03 mm and 0.01 mm, respectively. Additionally, each almond kernel was weighed using a scale (Mettler Toledo; Model - ML503T; Columbus, Ohio, USA) with a precision of 1 mg. The measurement process was repeated 10 times to obtain measurements for 1000 almond kernels.

### 2.3 Image processing algorithm

The image processing algorithm was applied by following the methods developed by Singh et al (2019), which is reiterated in brief below.

To ensure a fair contrast between the background and the brown kernels, the almond kernels were placed on clean dark grey surface. Along with almond kernels, for reference, an object (41 x 21 mm rectangular white paper strip) was also placed next to each image. Image of almond kernels was captured from the surface distance of approximately at 0.2 m using a cellphone camera (Apple, Cupertino, CA; Model: iPhone 7 Plus) with the camera flash turned off. To avoid the image distortion and shaking, the cellphone was mounted with a fixed tripod support. The acquired images were transferred to a personal computer for further image analysis.

The original image size obtained from the camera was 3553 x 2013 pixels. To ensure that the captured image had no objects other than the almond kernels and the reference paper strip, the original image was cropped. Afterwards, the cropped image was converted into a grayscale image by taking average of r, g and b values of the pixels (Eq 1). A binary image was derived by grayscaling the image using a single threshold value by comparing every pixel with a threshold value (*Th*_*pixel*_). A binary image comprises of either black (r = 0, g = 0, b = 0) or white pixels (r = 255, g = 255 and b = 255). A pixel with grayscale value below the threshold (*Th*_*pixel*_) was presumed to be background or black pixel and a pixel with grayscale value equal to or above the threshold was presumed to be representing the almond kernel or white pixel (Eq 2). *Th*_*pixel*_ value was based on the lighting and contrast between the background and the almond kernel. For every image, we manually adjusted the value of *Th*_*pixel*_ to cancel the background noise caused by dust or reflection while maintaining the integrity of the images of the almond kernels, which produced standardized binary images of the almond kernels. Furthermore, the recursive algorithm was used to identify each almond kernel from the image.

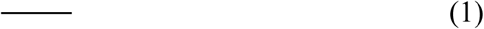

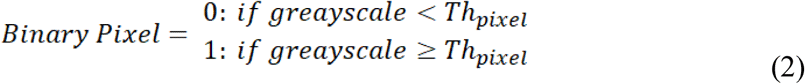

The almond images were processed using recursive algorithm which enables a function to to call itself from within. The recursive function tags every previously untagged neighboring white-colored pixel by visiting each pixel of white color (Arora et al., 2008) as shown in Fig 1. While scanning through every untagged adjoining pixel, the untagged neighboring white-pixels are tagged again. The tagging process ends when all the interlinked white-pixels are tagged. At the end of the function call, the algorithm identifies an almond kernel in the image by counting the number of tagged pixels. The recursive method scans through each white-pixel in a cluster of connected pixels regardless of the starting pixel, where a cluster of white-pixels symbolizes a single almond kernel in the binary image. In this case, any white-pixel, which was already tagged, is ignored by the recursive function. The algorithm continues to scan the binary image until all the white-pixels are tagged. At the end of the scanning process, every cluster is considered a separate object.

**Fig. 1.**
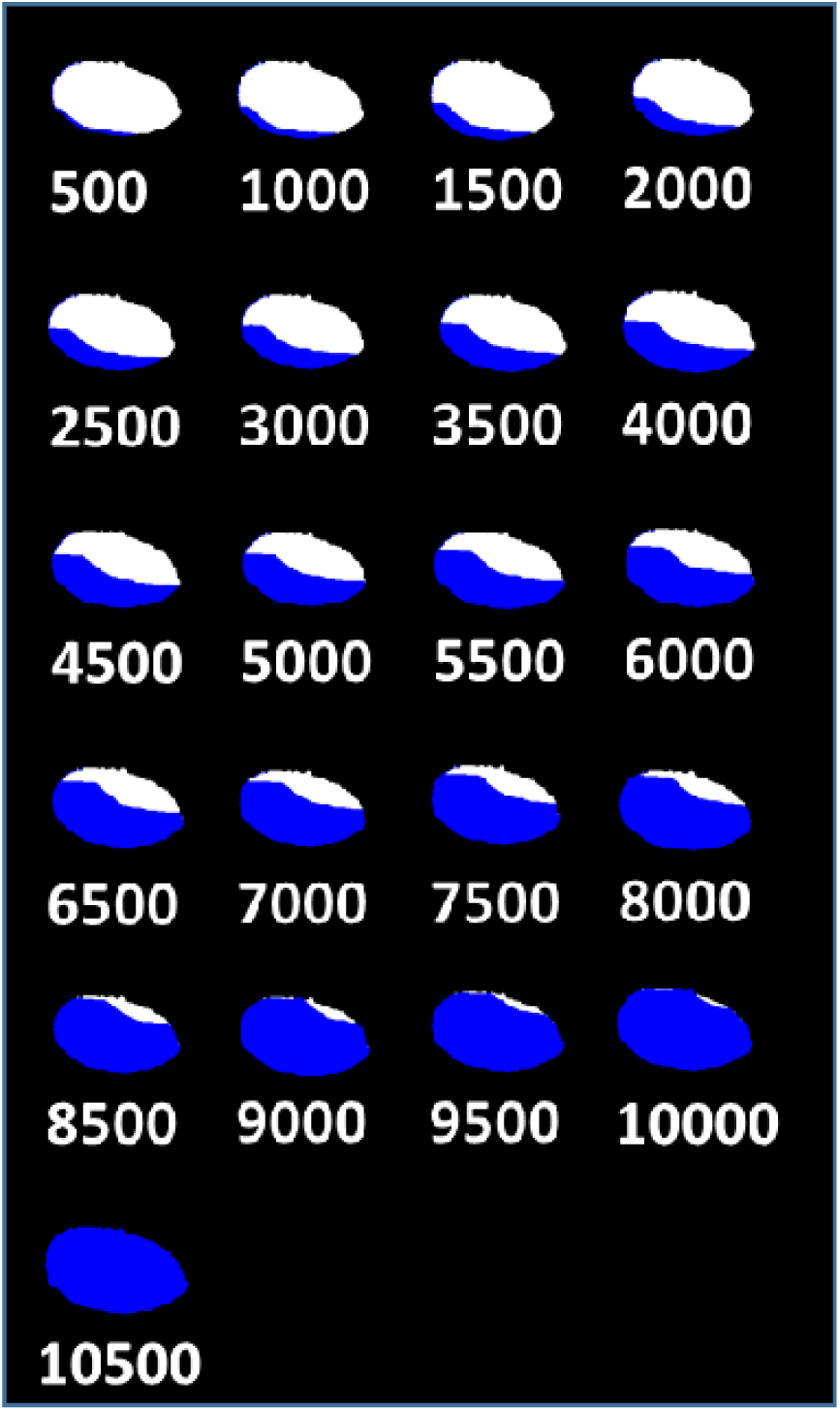
A representation of recursive algorithm tagging the white connected pixels of an almond kernel. For clarity, the tagged pixels are shaded blue. The number under each image symbolizes the number of recursive calls or number of white-pixels tagged. Once all the linked white-pixels are tagged, the recursive function ends.

The images are calibrated and converted from pixel domain to real world dimensions with the help of a white paper strip (41 x 21 mm) included in every image, as shown in Fig 2. By keeping the size of the white strip larger than any almond kernel, the code easily isolates the white strip as the largest cluster amongst the connected white-pixels and use it to calibrate the almond kernels. The spacing between every pixel from other pixels in the cluster was computed which helps calculate the kernel size. We assumed that the two pixels with the largest spacing were at the both ends along the length of the kernel. The length of the kernel 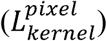 represents the distance between these two pixels in units of pixels. A green circle of diameter equal to the length of the pixel was drawn, with its center being the midpoint between the two end pixels. In addition, using Eq. 3, the length of the kernel 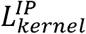 is converted from pixel to mm (Singh et al., 2019).

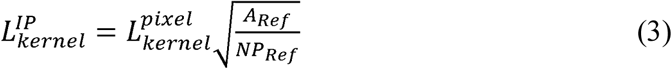

**Fig. 2.**
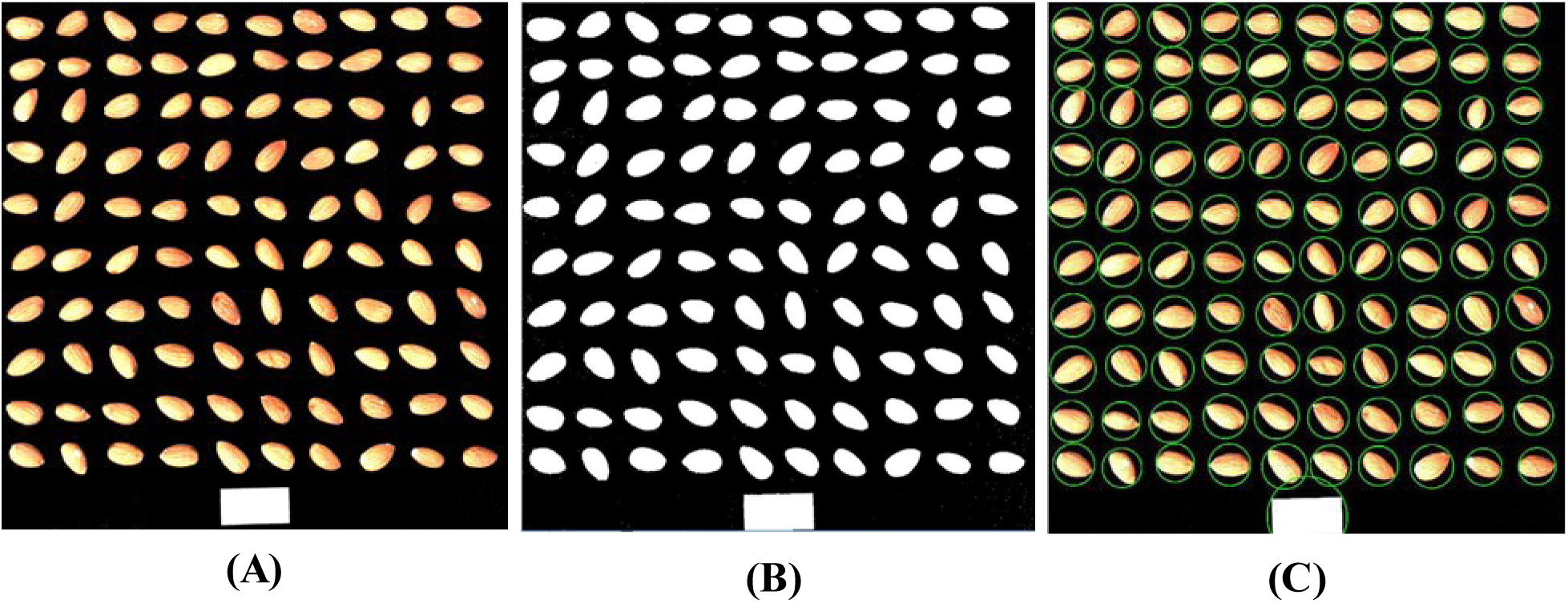
Phases for almond kernels identification. (A) original image as taken by the camera; (B) transformation of original image into binary image using thresholding process; and (C) identification of cluster of white pixels (almond kernels) using recursive function.

### 2.4 Machine learning

#### 2.4.1 Stacked Ensemble Model (SEM)

In this study, we have implemented a unique machine learning technique, where a collection of various ML models improved the prediction accuracy of the size and mass of the individual almond kernel from the image processing (Singh et al., 2019). Stack ensemble model (SEM) is a model which assembles different ML models in layers where the data flow forward from the input to the output layer.

An SEM usually has multiple layers where a data point contains an input feature vector (independent variables) and the output properties (response variables) corresponding to the input feature set (Fig 3). Based on the provided input feature vector to the ML models of the first layer, each ML model in the first layer estimates the output variable. The resulting estimate of individual ML model in the first layer is then combined together in the form of a *N*_*1*_ sized vector (*V*_*1*_), where *N*_*1*_ is the number of ML models in the first layer. Each ML model in the second layer utilizes the feature vector *V*_*1*_ as input. The ML models establish predictions for the size and mass output of the kernel in the second layer with the help of *N*_*1*_ ML models. Our SEM utilized only two layers, where the second layer was the output layer. To construct the SEM for this study, we used some of the most common ML models, such as Artificial Neural Network (ANN), Random Forest (RF), Support Vector Machine (SVM), Kernel Ridge Regression (KRR), and k-Nearest Neighbors (k-NN), as shown in Fig 3. This research implemented Standard ML libraries of the Scikit-Learn for python.

**Fig. 3.**
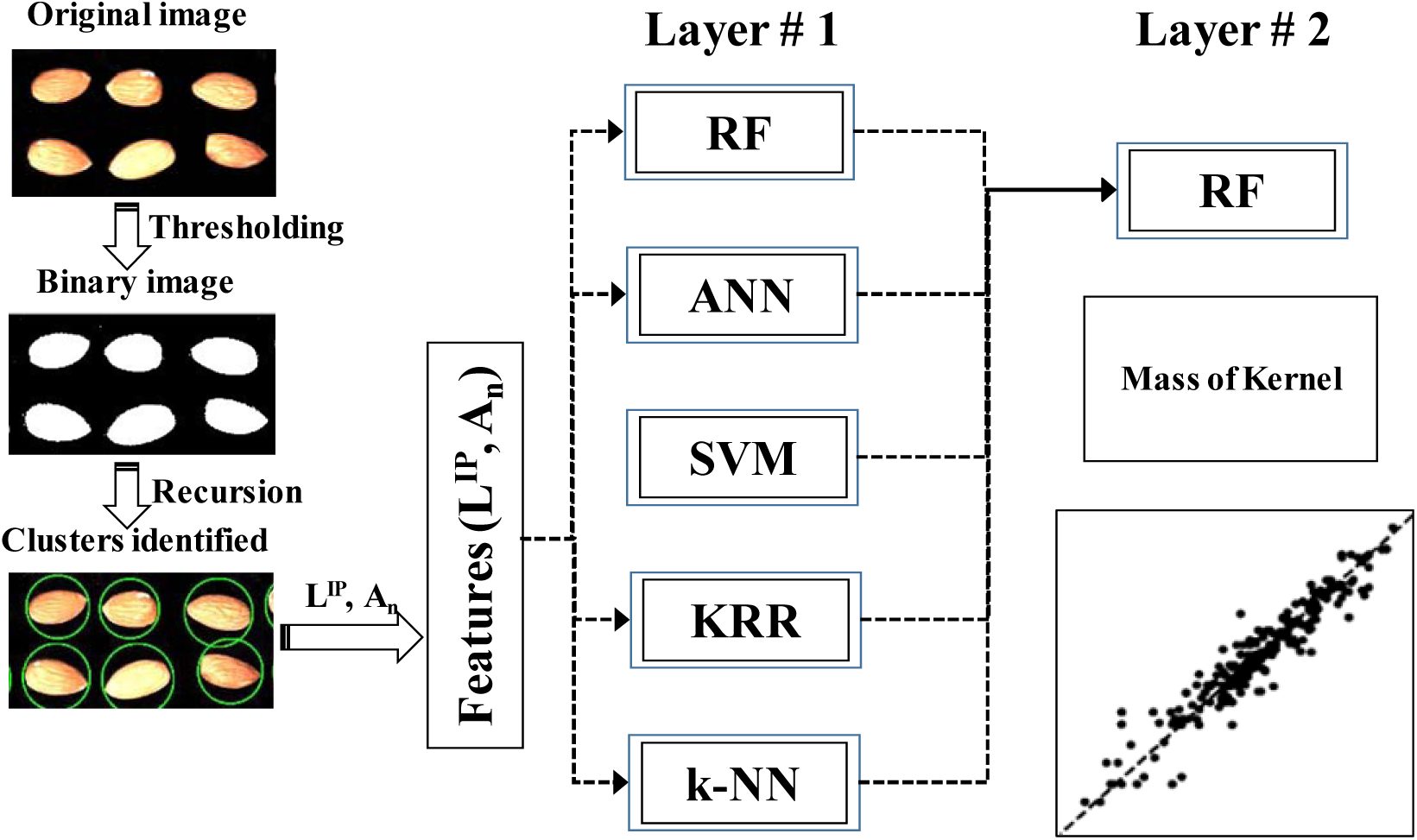
Process flow to predict the size and weight of the almond kernels

##### 2.4.1.1 Artificial Neural Network (ANN)

To construct predictive models, Artificial Neural Network (ANN) method is getting very popular in the science and engineering field. Several researchers have used ANN to classify the almond kernel (Ding and Gunasekaran., 1995; Reshadsedghi and Mahmoudi, 2013; Teimouri et al., 2016). There are many activation functions available and they get distinguished according to the threshold and shape of the filter function. The most common activation functions are ReLu, ELU, sigmoid, tanh and linear (Singh and Abbassi, 2018). We have used ReLu activation function for this study. An ANN model contains various nodes and layers of nodes as shown in Fig 4. A node is a computing unit that accepts an input signal from the previous layer’s node. After performing multiplication with weighting factors based on the input signal values, the summation of resulting value is fed through an activation function that filters and transforms the signal. Every node of the next layer receives the modified signal from the activation function.

**Fig. 4.**
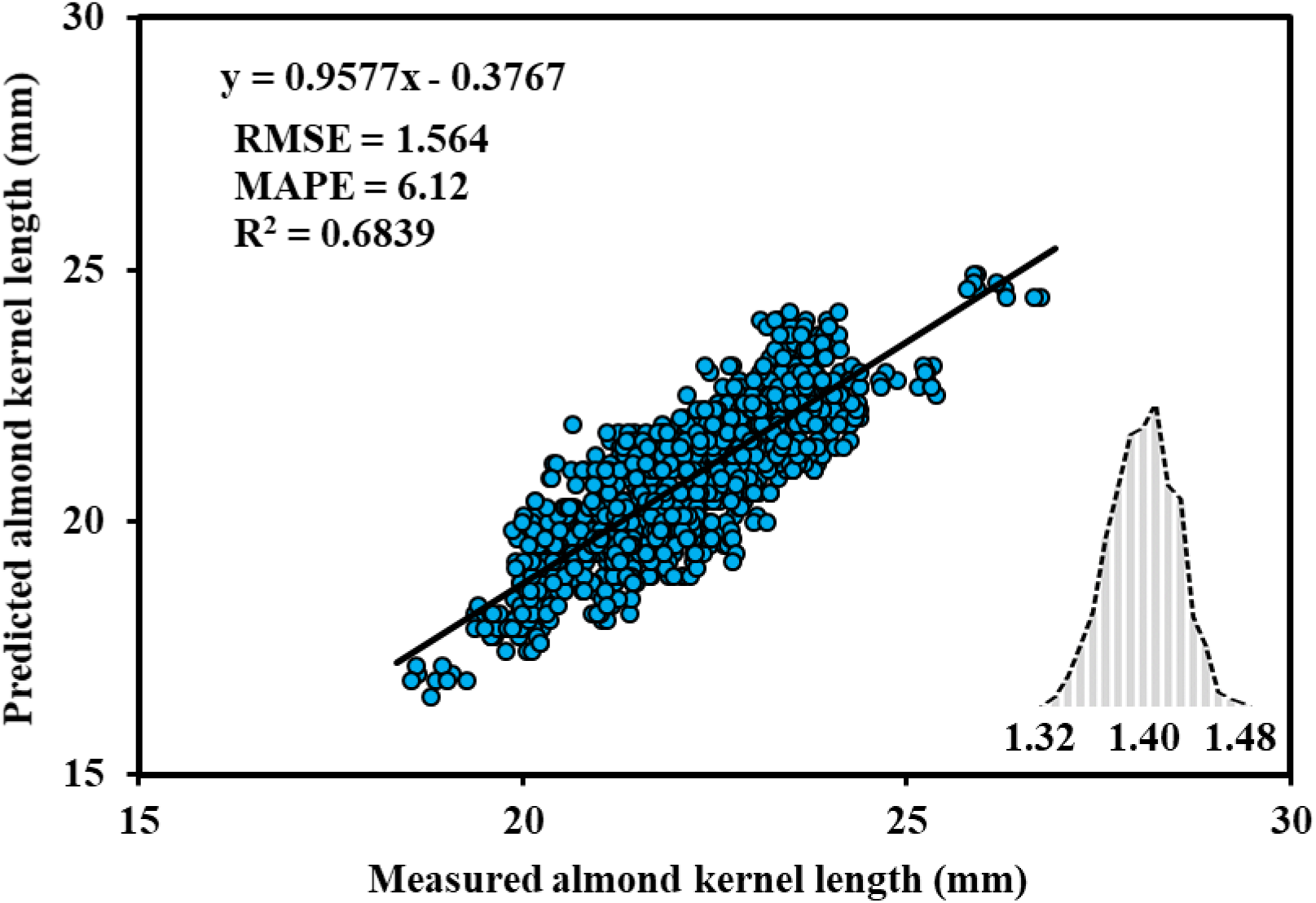
Measured kernel length vs predicted kernel length of almond

To minimize the error in estimation, an optimizer is used that tunes the weighting factors of every node throughout the training process.

##### 2.4.1.2 Random Forest (RF)

Random forest is a common and effectively managed machine-learning model, which is collaborative, comprises of various decision tree (DT) models (Farid et al., 2014; Quinlan, 1987; Silins et al., 2012; Yu et al., 2010). The final decision established by the RF model depends upon the voting or averaging of the underlying DT models. A DT logical model is generated by dividing and re-assembling the training data depending upon their similarity. Each node in a DT contains logical expression that further helps in deciding the coherent placement of a data point to the neighboring node situated in the left or right side of that particular node. *Root* is the initial node of the decision tree and the following splits are referred as *branches*. Leaves are the end nodes that indicates no data exists any further. Using a trained DT for prediction, the input data is fed through the root node, which then depending upon the logical expression flows through the branches and halts at leaf node. An average of linear regression of the training data points resulting from the leaf node is the output of a DT. The training data sets decide the shape and parameters of a DT. For training the DT models, multiple data sets are produced by bootstrapping the initial training data. Arbitrary picking data points from the original data set in such a way that a data point might get selected multiple times or not at all is called as bootstrapping. It is possible that from the original data set, duplicate points are selected or no points in a bootstrapped data set. The bootstrapping process generates unique data sets that ensures to have a forest of diverse decision trees. RF is one of the fastest ML model due to its plain and simple architecture of the original DT.

##### 2.4.1.3 Support Vector Regressor (SVR)

The standard Support Vector Machine (SVM) ML model has regressor variant called SVR which usually helps in classification application (Christianini and Taylor, 2000). SVM was applied to classify rice kernels, including head and broken rice kernels (Chen et al., 2012; Kaur and Singh, 2013). In n-dimensional feature space, it is relatively difficult to segregate the training data with a linear hyperplane, where ‘n’ is the number of features in the training data set. The fundamental algorithm of the SVR model converts the input features from original n-dimensional space to a higher dimensional space, fragmenting data sets into multiple homogeneous fragments using linear hyperplanes. The coefficients (*a*_*1*_, *a*_*2*_, *a*_*3*_, …, *a*_*n*_ and *b*) of the hyperplanes (Eq. 4) are optimized throughout the training process. Optimization maximizes the boundary (δ) between the training data points and the hyperplanes.

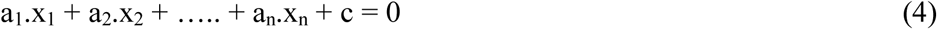

Converting the training data set from original n-dimensional feature space to a higher dimensional space is achieved through the kernel-trick method. In the converted space, the data points are linearly divisible with the help of the hyperplanes. The most commonly used Radial Basis Function (RBF) as described by Eq 5 is implemented in this study, where *k(x*_*i*_,*x*_*j*_) is the kernel function among two points *x*_*i*_ and *x*_*j*_ and *σ* symbolizes the kernel width. In this study, the value of *k*(*x*_*i*_,*x*_*j*_) signifies the resemblance amongst two almond grains. For two similar almond kernels, the value of *k*(*x*_*i*_,*x*_*j*_) → 1.

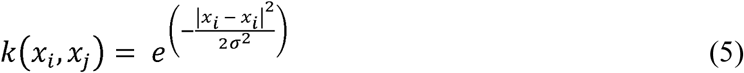

##### 2.4.1.4 Kernel Ridge Regression (KRR)

Similar to SVR, with the help of kernel-trick method, a Kernel Ridge Regression converts the training data set from original feature space to a higher dimensional space. In the higher dimensional space, the training data are divisible with linear hyperplanes. For example, the predicted length for almond kernel L(x_i_) with feature x_i_, can be denoted by the Eq 6.

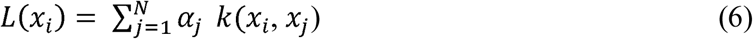

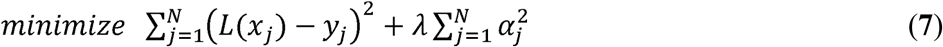

The objective function in Eq 7 is minimized to obtain vector α, where λ is a regularization penalty term and y_j_ is the measured length of the almond kernel. KRR is not extremely scalable and as the size of training data set grows, the computational speed suffers.

##### 2.4.1.5 k-Nearest Neighbors (kNN)

The k-nearest neighbors ML method predicts based on the average from the k nearest training data points to the input feature vector (Aurlien, 2017). In this study, three neighboring points (k=3) were used to predict the results. We used the Euclidean distance (Eq 8) to locate the three closest neighbors. Based on the input feature vector (*A*_*n*_ and 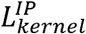), the algorithm computes its Euclidean distance from all the training data points in 2-dimensional domain. The mass of the almond kernel is predicted by averaging the masses of the three almond kernels closely matching its *A*_*n*_ and 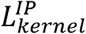 from the training data set. This is the most agile ML model due to its simplicity and accomplishes better results for evenly spread large data set in a featured space, especially in the region of enquiry. Since it stores all the training data points, it can be challenging with large training data set.

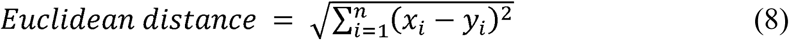

### 2.5 Performance of SEM

We evaluated the performance of our regression model using Root Mean Square Error (*RMSE*), Mean Absolute Percentage Error (*MAPE*) and correlation coefficient (*R*^*2*^) as shown in Eq 9-12. ‘x_i_’ and ‘y_i_’ represent the predicted and actual values, respectively, for ‘i = n’ number of predictions.

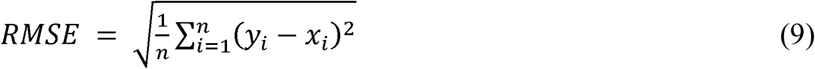

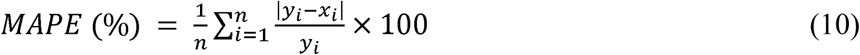

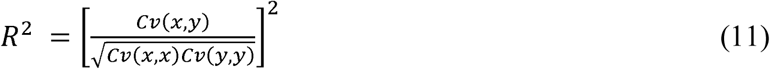

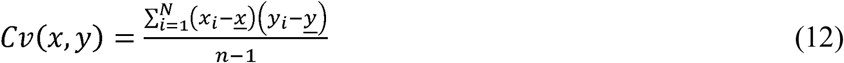

### 2.6 Uncertainty quantification of ML model (Bootstrapped statistics)

To understand the expected error in a prediction model, the residual errors of a ML model are typically quantified using single score indicator, such as RMSE and MAPE. However, these do not postulate the robustness and uncertainty of the ML prediction model. Therefore, in this study, the bootstrapping technique was used to generate several data sets and evaluate the average residual for each data set. The bootstrapping method randomly selects data points from the original data set *D*_*O*_ of size *n* and creates *N*_*BS*_ = 1000 new data sets, each of size n. The bootstrap can pick the same data points from the original data set multiple times or not pick at all. We used the SEM model to make the prediction for 1000 bootstrapped data set. The range of the uncertainty was calculated using the variance of the mean residuals for each data set. The accuracy and the robustness of the ML model were represented by the average and variance of the mean residuals of the bootstrapped data, respectively. A ML model with narrow distribution and lower average of the mean residuals was considered to be superior and desirable.

## 3 Results and Discussion

### 3.1 Almond kernel length estimation by image processing algorithm

We compared the measured length values 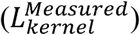 of individual kernels with the predicted values by the image processing method 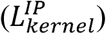 (Fig 4). The average measured kernel length of almond was 22.10 mm, whereas the corresponding length predicted by the image processing method was 22.79 mm with *R*^*2*^ = 0.68. The bootstrapped errors for 1000-kernel average length was found to be 1.40 ± 0.08 mm (Fig 8). The RMSE and MAPE values were 1.56 mm and 6.12%, respectively.

The differences between the 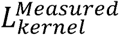 and 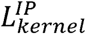 could be mainly due to measurement errors and image processing errors. The sources of image processing errors may be ascribed to calibration error, thresholding error, and image resolution. The calibration error can be caused by errors in measurements of the reference object dimensions which can result in the errors in almond kernel size estimation. Thresholding error can occur due to error based on threshold values during conversion of original image into the binary image during thresholding process. A pixel matching with the almond kernel may be documented as the background pixel (rgb = 0, 0, 0) and vice versa, which can lead to an under or over estimate of the number of pixels of a kernel. The resolution of the image is also a key factor attributing error. A higher resolution image has more pixels per almond kernel with smaller least count than a lower resolution image. The smaller kernels have fewer number of pixels with relatively smaller signal to noise ratio and prediction accuracy than the bigger kernels. Therefore, the use of a high image resolution camera can further improve the prediction accuracy and thus is recommended. In addition to a higher image processing error, the measurement errors are also higher in smaller object.

### 3.2 Almond kernel mass estimation by image processing algorithm

We hypothesize that the two major dimensions of the kernel are represented by the 2D projected image of an almond kernel which can be used as a variable to predict the mass of the almond kernel. The rectangular white strip with the known dimensions in the image was used as a standardized reference. The pixel area (number of pixels) representing an almond kernel was divided by the pixel area of the reference to obtain a standardized value and then multiplied with a constant *C*_*n*_ = 3.373 to obtain a normalized pixel area value (*A*_*n*_) for each almond kernel, as shown in Eq 13. The maximum value of *A*_*n*_ represented the largest almond kernel.

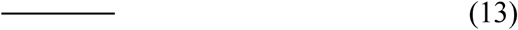

The measured mass of the individual almond kernel was correlated with its *A*_*n*_. The average measured mass of the almond kernel was 1.10 g. The *R*^*2*^ values (0.34) indicate a poor correlation between the *A*_*n*_ and the measured mass of the almond kernels as shown in Table 1 and Fig 5. The bootstrapped errors for kernel average mass was 0.28 ± 0.01 g. The RMSE and MAPE values were 0.101 g and 7.52%, respectively.

**Fig. 5.**
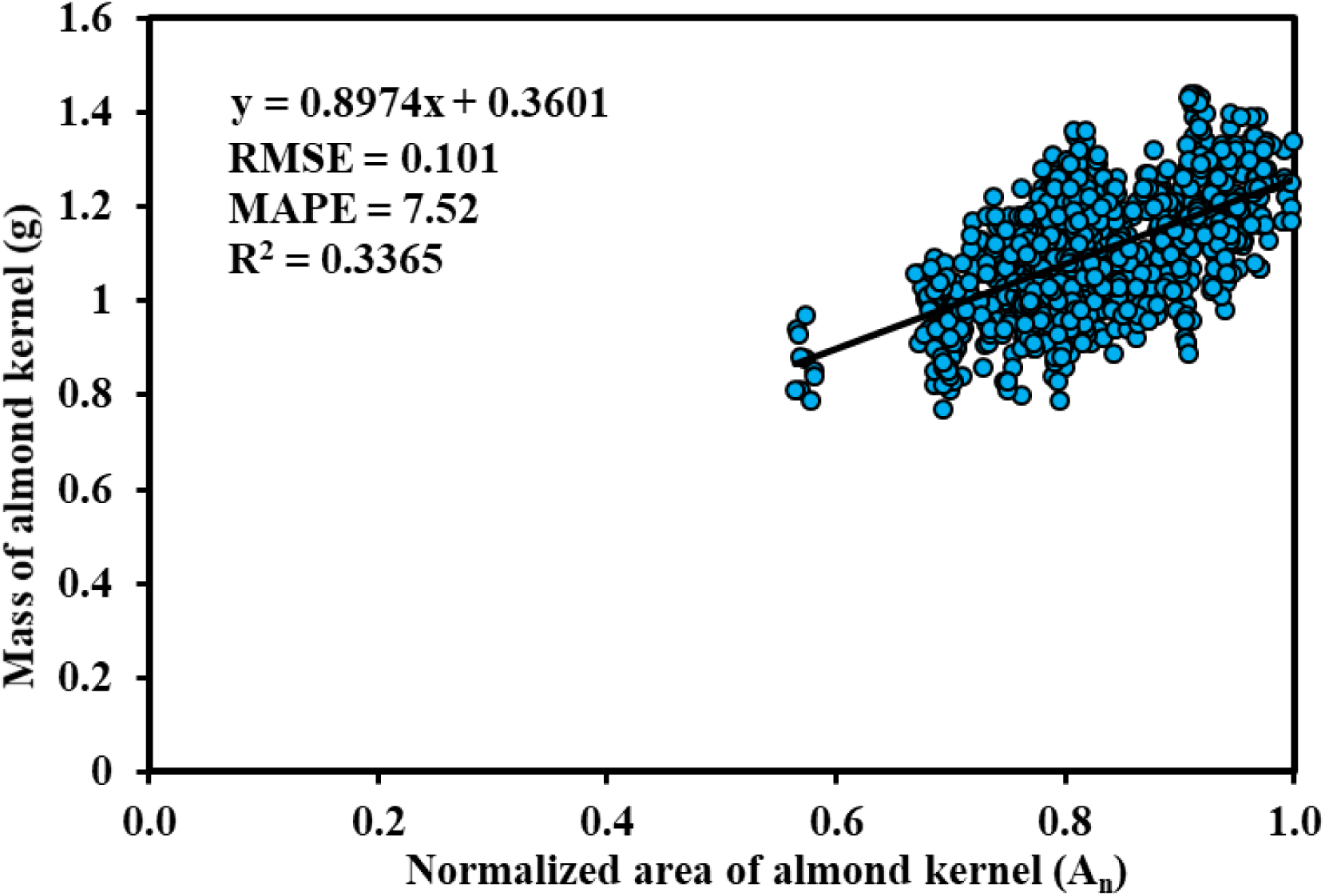
Normalized pixel area (A_n_) vs measured mass of almond

The higher errors and lower prediction accuracy could be attributed to the almond kernels being typically wider in one of its two radial directions (width and thickness), of which, the smaller breadth was ignored in this study. Since, the width and thickness of the almond kernels could be a key factor contributing to the weight-area prediction correlation, overlooking the thickness could possibly result in higher RSME and MAPE values. The prediction errors could mainly be accredited to the measurement, calibration, and thresholding errors, as well as image resolution. In addition, the error in the measurement of the reference object (rectangular strip) dimensions can propagate to the errors in almond kernel size estimation, ultimately reflecting in the correlation between *A*_*n*_ and its corresponding measured mass. As stated earlier, during the thresholding process, a pixel associated with the almond kernel may be acknowledged as the background pixel (rgb=0,0,0), leading to an underestimation of the number of pixels accompanying the image of almond kernel. On the other hand, some background pixels around the almond image could be identified as white (almond image) pixels and included in the cluster of almond image pixels, resulting in an overestimation of the *A*_*n*_.

Moreover, the image resolution also contributes to the overall error. A high-resolution image contains a higher number of pixels and a smaller value of the least count, which leads to a higher accuracy and finer details in acquiring the size and shape of the almond kernel. On the other hand, an image with lower resolution of the kernels contains fewer coarser pixels, causing either under- or over-estimation of the value of 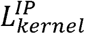 and *A*_*n*_. In order to explore the effect of image resolution, a test case with four almond kernels was studied and their 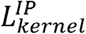 and *A*_*n*_ values were compared at different image resolutions (10% and 100%) as shown in Fig 6. We performed the coarsening operation of the original image using an image processing tool GIMP and a linear scaling method. The relative difference (RD) between quantities calculated using the lower resolution images and the original image (100% resolution) converged at around 75% resolution. The number of pixels representing the almond kernel in the original image is 9,482 pixels and for 75% resolution image is 7,112 pixels. It could be inferred that the image processing error due to resolution is negligible if the almond kernel in the image is represented by approximately 7,200 pixels or greater. We calculated the percentage image resolutions with respect to the original image of the almond kernel (Eq 14-15). The Fig 6 demonstrates that our image setting was enough to minimize the error due to image resolution.

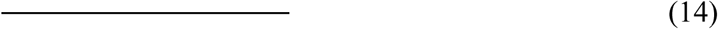

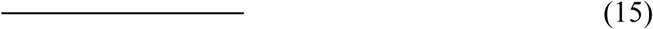

**Fig. 6.**
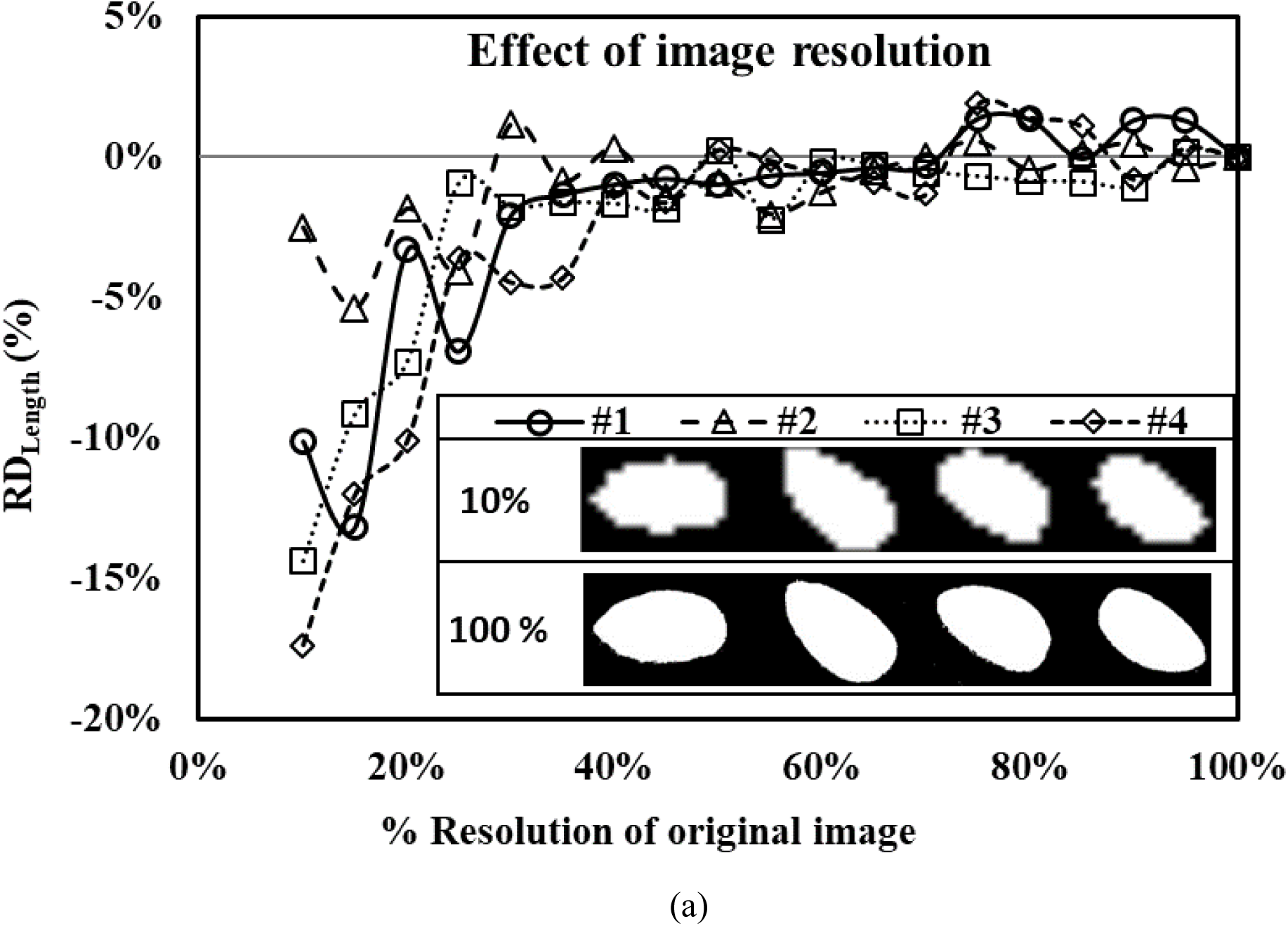
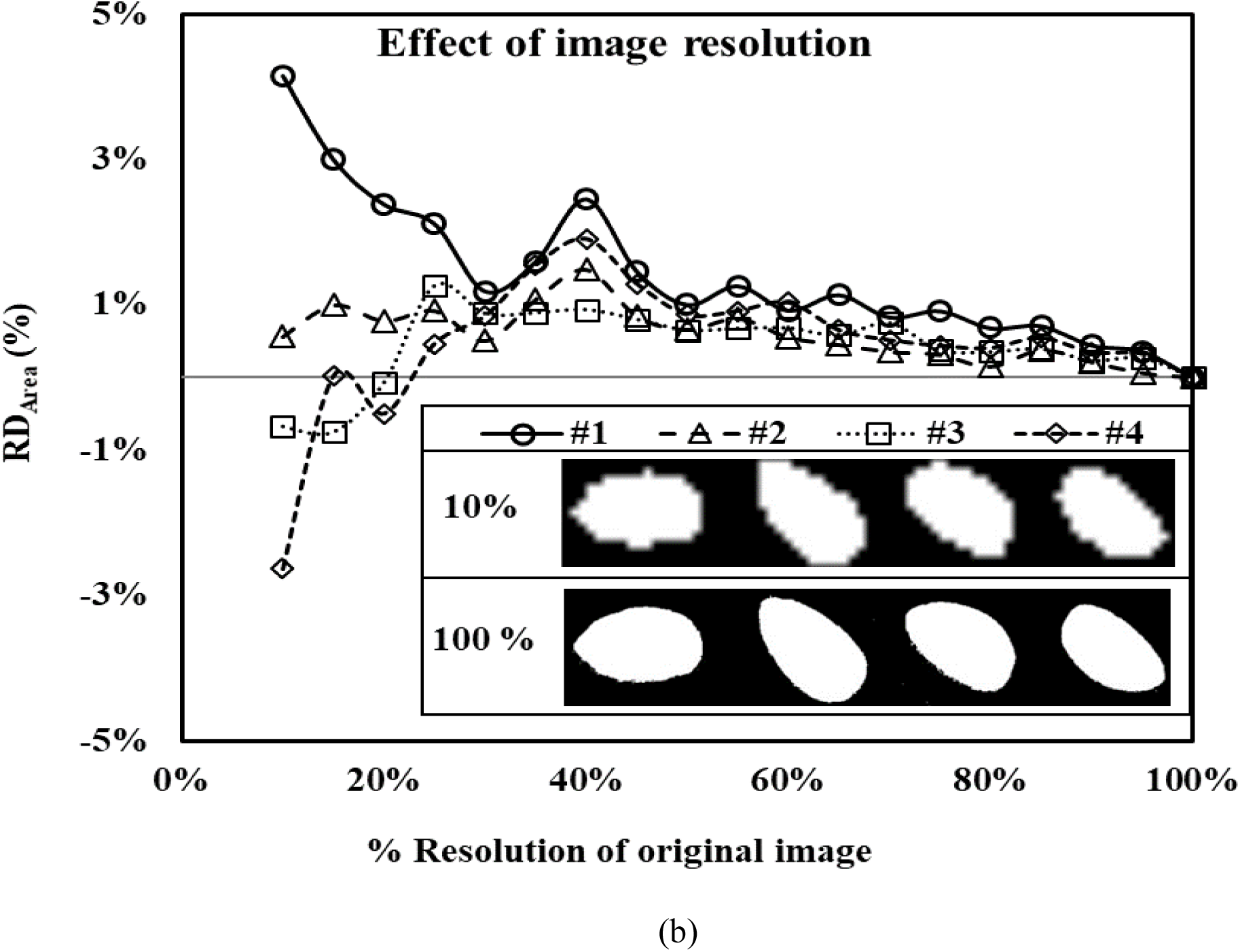
Relative difference due to the effect of image resolution on the calculated values of (a) and (b) *A*_*n*_ value for four random almond kernels. The insat images show four randomly selected almond kernels at 10 % and 100% resolutions of the original image.

The prediction estimates for almond mass and model accuracy were further analyzed using SEM. The average mass of almonds as predicted by SEM was 1.107 g. The SEM model provided an improved estimation for the mass of the almond kernels compared to the linear least-square fit model. The prediction accuracy indicators (RMSE, MAPE and R^2^) obtained from SEM models are shown in Table 2 and Figure 7, which indicated the progressive improvement of the accuracy with the layers of ML models in the SEM model. For example, the RMSE value reduced from 0.070 g in RF: Layer 1 to 0.060 in RF: Layer 2 in the SEM model. Similar trends were noticed for MAPE and R^2^ where MAPE improved from 4.52% to 3.63% and R^2^ from 0.651 to 0.74 moving from RF: Layer 1 to RF: Layer 2 in the SEM model, respectively. The uncertainty quantification of residuals (mean error ± 95% limit) also indicated progressive accuracy improvements with progressive layers of ML models in the SEM model. For example, the uncertainty quantification of residuals improved from 0.48 ± 0.05 g in RF: Layer 1 to 0.38 ± 0.05 g in the RF: Layer 2 of the SEM model.

**Table 2.** Performance indicators for software predictions and ML models for the mass of the almond kernels.

| Software Prediction Stats |  |  | SEM Predictions Stats |  |  |  |  |
| --- | --- | --- | --- | --- | --- | --- | --- |
| RMSE (g) | MAPE (%) | R<sup>2</sup> |  | RMS E (g) | MAPE (%) | R<sup>2</sup> | Uncertainty quantification of residuals (Mean error ± 95% limit) |
| 0.101 | 7.52 | 0.336 | Layer 1: RF | 0.07 | 4.52 | 0.0651 | 0.48 ± 0.05 |
|  |  |  | Layer 1: ANN | 0.096 | 6.79 | 0.339 | 0.74 ± 0.06 |
|  |  |  | Layer 1: SVM | 0.091 | 6.48 | 0.52 | 0.70 ± 0.06 |
|  |  |  | Layer 1: KRR | 0.082 | 5.24 | 0.52 | 0.56 ± 0.06 |
|  |  |  | Layer 1: k-NN | 0.091 | 6.6 | 0.0407 | 0.70 ± 0.06 |
|  |  |  | Layer 2: RF | 0.06 | 3.63 | 0.743 | 0.38 ± 0.05 |

**Fig. 7.**
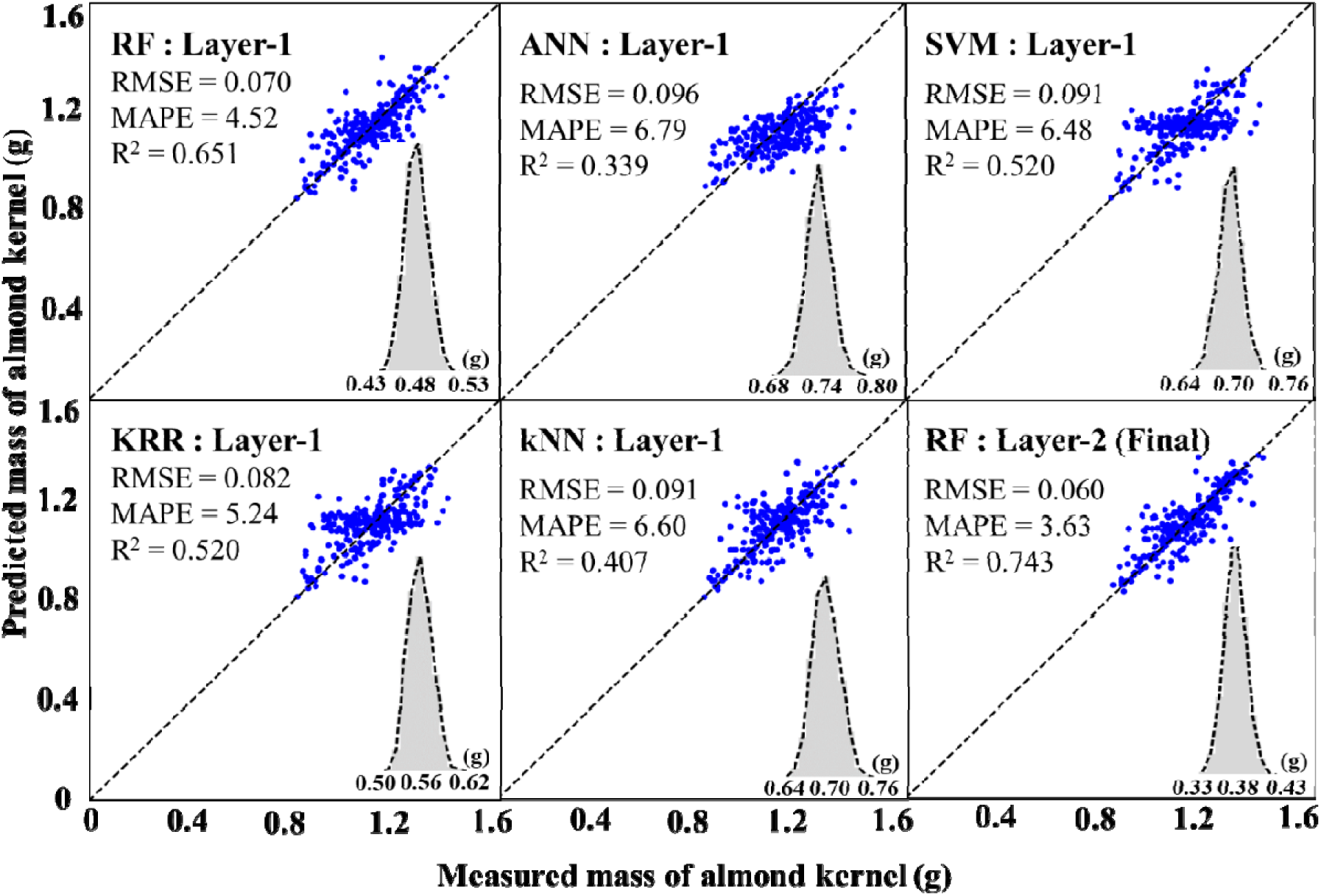
SEM Predicted mass vs measured mass of almond kernels

## 4 Conclusions

An application of image processing and machine learning was successfully demonstrated to accurately measure the size and mass of individual almond kernels with high confidence level. The size and area of the individual almond kernels were obtained in the form of pixels using image-processing algorithm. Further, the pixel information of individual almond kernel was converted into feature vectors, which represented their size and area and predicted their lengths and weights. Several ML models were stacked to create an SEM, which improved the prediction accuracy moving from one layer to the next. The final prediction accuracy obtained from the SEM was greater than the ML model or least square error fit, signifying the supremacy of stacking the ML models. We also studied the effect of image resolution on the image processing error and found that the errors converged with the improved resolution with approximately more than 7200 pixels.

We used a California-grown almond cultivar (classification – Nonpareil) to evaluate our methodology. The average measured length of the 100 kernels was 22.10 mm. Using our image processing algorithm, we estimated the average length of the 100 kernels to be 22.79 mm and bootstrapped residuals with 95% confidence interval of 1.40 ± 0.08 mm.

Using SEM, we estimated the average mass of the individual almond kernels which was 1.107 g. The bootstrapped residuals with 95% confidence level was 0.38 ± 0.05 g. The low values of mean errors and narrow ranges of the 95% confidence intervals specified the robustness and accuracy of our methodology. This study can have important applications in industrial almond processing, which has capability to replace the manual and tedious methods typically applied for kernel size and weight measurements, saving enormous amount of time and effort. Moreover, the use of the developed method in image analyzers can offer an objective assessment of the industrial almond yield.

